# Antimicrobial Efficacy and Safety of a Novel Gas Plasma-Activated Catheter Lock Solution

**DOI:** 10.1101/302919

**Authors:** Sudhir Bhatt, Poonam Mehta, Chen Chen, Dayle A. Daines, Leonard A. Mermel, Hai-Lan Chen, Michael G. Kong

## Abstract

Antimicrobial lock solutions are important for prevention of microbial colonization and infection of long-term central venous catheters. We investigated the efficacy and safety of a novel antibiotic-free lock solution formed from gas plasma-activated disinfectant (PAD). Using a luminal biofilm model, viable cells of methicillin-resistant *Staphylococcus aureus, Staphylococcus epidermidis, Pseudomonas aeruginosa*, and *Candida albicans* in mature biofilms were reduced by 6 - 8 orders of magnitude with a PAD lock for 60 minutes. Subsequent 24-hour incubation of PAD-treated samples resulted in no detectable regrowth of viable bacteria or fungi. As a comparison, the use of a minocycline/EDTA/ethanol lock solution for 60 minutes led to regrowth of bacteria and fungi, up to 10^7^ - 10^9^ CFU/ml, in 24 hours. The PAD lock solution had minimal impact on human umbilical vein endothelial cell viability, whereas the minocycline/EDTA/ethanol solution elicited cell death in nearly half of human endothelial cells. Additionally, PAD treatment caused little topological change to catheter materials. In conclusion, PAD represents a novel antibiotic-free, non-cytotoxic lock solution that elicits rapid and broad-spectrum eradication of biofilm-laden microbes and which shows promise for the prevention and treatment of intravascular catheter infections.

## INTRODUCTION

Central venous catheters (CVCs) provide long-term access to medication and total parenteral nutrition for cancer, hemodialysis, short-gut, and transplant patients. The majority of bloodstream infections in long-term CVCs (>10 days) is associated with intraluminal microbial colonization (1,2). Current management guidelines for catheter-related bloodstream infections (CRBSIs) recommend use of antimicrobial lock therapy (ALT) for catheter salvage (3). Even with antibiotics at concentrations 1,000-fold above the systemic therapeutic dose, current ALT may be ineffective in eradicating mature bacterial and fungal biofilms (4-7). In addition, leakage of antimicrobial lock solutions into the bloodstream has been implicated in systemic toxicity in patients with long-term CVCs, potential for increased biofilm formation, as well as adverse effects on the catheter integrity and intraluminal precipitation (5-6,8-9).

After three decades of optimizing their utility in catheter lock solutions, reliance on current antibiotics is fundamentally challenged by the limited scope to improve the trade-off between efficacy and toxicity (6,7,10) and by the potential risk of antimicrobial resistance (11). Inspired by the way that endogenous reactive oxygen species (ROS) are released by immune cells of a mammalian host to inactivate invading bacteria, extensive studies have shown that exogenous reactive oxygen and nitrogen species (RONS) and other antimicrobial effectors (e.g. transient charges) generated by gas plasmas effect rapid inactivation of bacteria and fungi (12-14). In addition, RONS from gas plasmas can be designed to be selective against microbes with little harm to mammalian host (15,16).

We recently developed a novel gas plasma-activated disinfectant (PAD) as a novel catheter lock solution. The aim of the current study was to determine the efficacy of PAD against bacteria and fungi in a catheter biofilm model and to assess the effect of PAD on primary human umbilical vein endothelial cells (HUVEC) as an *in vitro* model of blood vessel endothelium. In doing so, we compared the PAD to a novel antibiotic-antiseptic lock solution.

## MATERIALS AND METHODS

### Catheter lock solutions

Minocycline hydrochloride (3 mg/ml) and EDTA (30 mg/ml) were mixed in 25% ethanol (M-EDTA-25E), as a comparator lock solution (17). PAD was formed by treating 5 ml of normal saline (NaCl) for 2 minutes with a room-temperature gas plasma system at 3.2 W in ambient air (18, Figure 1). Untreated saline was used as a control.

**Figure 1.**
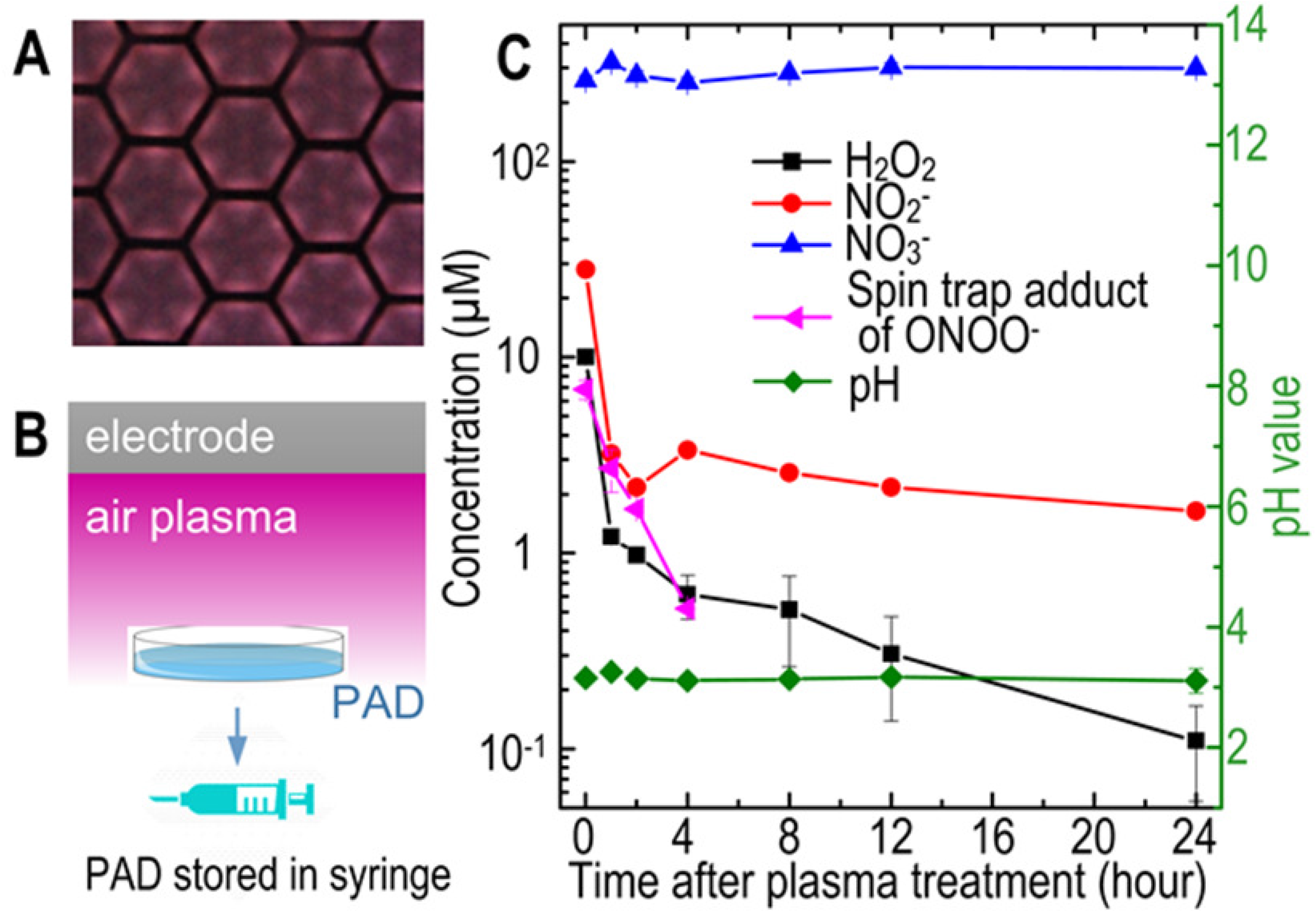
(A) End view of an atmospheric air plasma generated on the surface of its hexagon-shaped mesh electrode on a glass slide (in black) with each hexagon metal rim confining one surface discharge plasma (in purple). The air plasma was sustained at a peak-to-peak voltage of 9.4 kV at 23 kHz and a dissipated power density of 3.2 W. (B) A side view schematic of the surface plasma being used to treat 5 ml normal saline in a petri dish downstream from the electrode for 2 min. The resulting PAD was then stored in a sterile syringe. (C) Concentrations of H_2_O_2_, NO_2_^-^, NO_3_^-^, and spin-trap adducts of ONOO^-^ as well as pH of PAD at 37°C as a function of time after plasma activation of saline.

### Microorganisms and luminal biofilm model

MRSA BAA-1707, *P. aeruginosa* BAA-47 and *C. albicans* 14053 (ATCC, Manassas, VA) were used to test antimicrobial efficacy of PAD. Additionally, PAD was tested with clinical isolates of MRSA (SO385), *S. epidermidis* (M0881) and *C. albicans* (BEI, Manassas, VA), as well as a laboratory control strain of *P. aeruginosa* 27853 (ATCC, Manassas, VA). *MRSA* and S. *epidermidis* were grown on Luria-Bertani (LB) agar, *P. aeruginosa* on Brain Heart Infusion (BHI) agar, and *C. albicans* on Yeast-Malt (YM) agar. A single colony was inoculated into 10 ml of appropriate broth, incubated overnight at 37°C while shaking at 160 rpm, harvested at mid-logarithmic phase growth by centrifugation (500 x g, 5 min), then washed twice with 1xPBS. The inoculum concentration was adjusted to 1.0 - 2.0 x10^7^ CFU/ml by broth dilution.

We modified a luminal biofilm model (4,19) using sterile silicone tubing of 1.58 mm ID, 3.175 mm OD (NewAge, Southampton, PA). Silicone tubing segments, 305 mm in length, were sterilized and inoculated with approximately 600 μl of a prepared microbial culture using a sterile syringe. Each inoculated catheter was sealed with a sterile, tight-fitting PTFE plug, placed in a sterile Petri dish, and incubated without shaking for 24 or 48 hours at 37°C. Inoculated segments were gently flushed with normal saline, removing non-adherent microorganisms and leaving intact biofilm.

### Eradication and regrowth assays

In the eradication assay, 600 μl of a test lock solution was slowly injected into the lumen of each inoculated segment using a syringe. Segments were then incubated at 37°C for 15 - 360 minutes. After incubation, the external surface of each segment was thoroughly swabbed with 70% (v/v) ethanol and allowed to air dry to ensure that only luminal microbes were recovered. Luminal contents were removed with a gentle saline flush, each segment was then cut into three segments of 101.6 mm and submerged in 2 ml of 0.1 M glycine buffer (pH 7.0). Each of these segments was then vortexed for 1 minute, sonicated at 40 kHz for 1 minute in a room temperature water bath, then vortexed for an additional minute. Luminal contents were then serially diluted and colony counts were enumerated using a plate counter (detection limit 10 CFU/ml).

To evaluate microbial regrowth, catheter segments from each eradication treatment were placed in a sterile 15 mil Falcon tube filled with 2 ml of appropriate broth and incubated overnight with shaking at 37°C. Following incubation, segments were vortexed for 1 minute, then sonicated for 1 minute. A 200 μl aliquot of sonicated broth was used for serial dilution and plating. Plates were incubated overnight at 37°C before colony enumeration.

### Biofilm Dispersal Assay

Glass chamber slides (Thermo Fisher Scientific, Fair Lawn, NJ) were inoculated with 500 ul of microbial cultures at 1.0 - 2.0 x10^7^ CFU/ml and then incubated for 24 hours to form biofilm on the surface of the slides. To remove planktonic cells, the medium was aspirated off and the biomass washed three times with 1 ml of 1x PBS. Following gentle washing, biofilms were treated with 200 μL of lock solution for 60 minutes. Before and after exposure to the lock solution, various wells of the inoculated slides were stained with crystal violet (BD, Franklin Lakes, NJ) for imaging and quantification of any adherent material (20). Adherent material on the glass slides was quantified by means of optical absorption at 590 nm.

### Cytotoxicity Assay

To assess for cytotoxicity of inadvertently spilled test lock solutions, we used HUVEC (Lonza, Walkersville, MD) with the MTT (3-(4,5-Dimethylthiazol-2-yl)-2,5-Diphenyltetrazolium Bromide) assay using the Vybrant^®^ MTT Cell Proliferation Assay Kit (Invitrogen, CA, USA) according to the manufacturer’s instruction. Briefly, HUVECs were seeded in endothelial basal medium (Lonza, Walkersville, MD) at 200 μl/well in 96-well plates at a density of 1.0 × 10^5^ cells/ml one day prior to a cytotoxicity test. To determine the volume of test lock solution to treat HUVECs, we used the algorithm that the maximum lock solution escaping from a CVC is approximately 0.5 ml per event (21) and this solution is distributed in the amount of blood pumped in 1 - 2 heart beats (22), which is approximately 100 ml in a healthy adult. This corresponds to a lock solution-blood ratio of up to approximately 0.5% per spill. Given this, each test lock solution was mixed at 0.1 - 0.5% of the cell medium in each HUVEC-containing well of the 96-well plates. Untreated HUVECs served as controls.

### Measurements of Reactive Species

Long-lived RONS in PAD at 37°C were measured with a microplate reader (FLUOStar, BMG Labtech, NC, USA) with the Amplex Red assay kit (Thermo Fisher Scientific, MA, USA) for H_2_O_2_ and the Griess Reagent assay (Cayman Chemical Co, MI, USA) for nitrite (NO_2_^-^) and nitrate (NO_3_^-^). Similarly, short-lived RONS were measured at 37°C with an electron spin resonance (ESR) spectrometer (EMX+, Bruker, Germany) and spin traps (18). Specifically, DMPO (5,5-Dimethyl-1-Pyrroline N-oxide) was used for trapping hydroxyl radicals (^·^OH), and DTCS (diethyldithiocarbamate) and MGD (N-methyl-D-glucamine dithiocarbamate) for trapping nitric oxide (NO^·^; all from Dojindo Laboratories, Kumamoto, Japan). Superoxide (O_2_^·-^) and peroxynitrite (ONOO^-^) were trapped using TEMPONE-H (1-hydroxy-2,2,6,6-tetramethyl-4-oxo-piperidine hydrochloride, from Enzo Biochem, NY, USA). The ESR detection limit was approximately 100 nM.

### Surface characterization

Surface topology of the inner wall of silicone catheter tubing was examined using a JSM-6060LV scanning electron microscopy (SEM; JEOL, Japan). To mimic prolonged exposure, tubing segments were locked with 600 μL of PAD for 2 hours at 37°C. Then, lock solutions were replenished with 600 μL of freshly prepared PAD for another 2 hours at 37°C. This was repeated consecutively 5 times in a day. After the last 2-hour treatment, test segments were locked with freshly prepared PAD overnight (~ 14 hours) at 37°C. The treatment procedure lasted for 7 days. As controls, segments of silicone tubing were locked with normal saline with the same procedure for lock solution replacement and untreated segments were stored at 37°C, both for 7 days. Following the treatment, tubing segments were cut open and the inner surface of each segment was gold-coated and examined with SEM at a working voltage of 15 kV.

### Statistical Analysis

All test conditions were studied in at least three independent experiments. Data are presented as mean ± SD. The Student’s *t* test was used to determine significance between data points.

## RESULTS

### Transient reactive oxygen and nitrogen species

Diverse reactive oxygen and nitrogen species in PAD was observed with all long-lived RONS at a low concentration, with peak values at 10 - 300 μM (Figure 1). The peak concentration of a given reactive species in PAD was markedly below the minimum inhibitory concentration (MIC) of the species when acting alone. For example, the MICs of H_2_O_2_ and ONOO^-^ are both 1 - 10 mM for *E. coli* (23, 24) and these are approximately 3 orders of magnitude above the peak concentrations of H_2_O_2_ and spin-trap adducts of H_2_O_2_ (10 μM) and ONOO^-^ (7 μM) found in PAD (Figure 1). Figure 1 shows half-lives of 30 - 45 minutes for H_2_O_2_, NO_2_^-^ and ONOO^-^ as well as plasma-induced acidification (pH = 3) with which the main reactive chlorine species was hypochlorous acid (HOCl) below 100 nM in air plasma-activated saline of greater than 1 mm in thickness (25), 3 orders of magnitude below its MIC of approximately 12.5 μM against S. *aureus* (26). Peak concentrations of NO_2_^-^ (30 μM) and NO_3_^-^ (300 μM) were much lower than their MICs (>0.5 mM for NO_2_^-^ and >10 mM for NO_3_^-^) (27) and cytotoxicity dose (>18 mM for NO_3_^-^) (28). Together, antimicrobial reactive species in PAD were at least 3 orders of magnitude below their individual MICs and were transient; with H_2_O_2_, NO_2_^-^ and ONOO^-^ half-lives of 30 - 45 minutes. Fundamentally distinct from antibiotics, PAD is characterized by chemical diversity of many RONS, each at low concentrations and decaying rapidly within one hour.

### Antimicrobial and Antibiofilm Efficacy

From an initial microbial load of 10^7^-10^8^ CFU/ml, the PAD lock eradicated 24-hour luminal biofilms of *MRSA* (BAA-1707) and *P. aeruginosa* (BAA-47) within 30 minutes and *C. albicans* 14053 within 45 minutes (Table 1A). Complete suppression of regrowth in the segment lumens was confirmed at 24 hours after an extended lock of 45 minutes for bacteria and 60 minutes for *C. albicans* (Table 1B). However, the M-EDTA-25E treatment left a residual bacterial population of 3.0 - 3.5 log_10_ CFU/ml with 30-minute incubation and a residual fungal population of 3.5 log_10_ CFU/ml with 60-minute incubation (Table 1A). Furthermore, 24-hour incubation of segment lumens treated with M-EDTA-25E lock solution for 60 minutes led to recovery of *P. aeruginosa* and *C. albicans*, both to approximately 4.4 log_10_ CFU/ml (Table 1B). Against clinical isolates, PAD lock for 60 minutes led to a 6-8 log reduction of all bacterial and fungal test microorganisms (Table 2A) and complete suppression of regrowth (Table 2B). The same 60-minute exposure to M-EDTA-25E lock solution resulted in approximately 2.8 log_10_ CFU/ml of MRSA and S. *epidermidis* and 4.3 log_10_ CFU/ml of *C. albicans* in segment lumens (Table 2A). Microbial recovery with M-EDTA-25E reached the initial luminal inoculum of 7 - 9 log_10_ CFU/ml (Table 2B).

**Table 1.** Survivors (A) and regrowth (B) of ATCC-derived isolates following PAD treatment

| Lock Solution | Lock time (min) | Survivors ($\log_{10}$ CFU/ml ± SD ) | | |
| --- | --- | --- | --- | --- |
|  |  | *MRSA* BAA-1707<sup>a</sup> | *P. aeruginosa* BAA-47<sup>a</sup> | *C. albicans* 14053<sup>a</sup> |
| Untreated | 0 | 7.97 ± 0.04 | 7.95 ± 0.03 | 6.92 ± 0.23 |
| Saline | 60 | 7.58 ± 0.07 | 7.70 ± 0.07 | 6.56 ± 0.09 |
| M-EDTA-25E | 15 | 5.30 ± 0.01 | 5.44 ± 0.04 | 4.80 ± 0.36 |
| | 30 | 3.46 ± 0.10 | 2.97 ± 0.17 | 4.08 ± 0.18 |
| | 45 | <1 | 1.76 ± 0.44 | 3.87 ± 0.15 |
| | 60 | <1 | <1 | 3.47 ± 0.23 |
| PAD | 15 | 4.91 ± 0.11 | 3.92 ± 0.43 | 4.90 ± 0.35 |
| | 30 | <1 | <1 | 3.21 ± 0.26 |
|  | 45 | <1 | <1 | <1 |
|  | 60 | <1 | <1 | <1 |

(B)
| Lock Solution | Lock time (min) | Regrowth ($\log_{10}$ CFU/ml ± SD) | | |
| --- | --- | --- | --- | --- |
|  |  | *MRSA* BAA-1707<sup>a</sup> | *P. aeruginosa* BAA-47<sup>a</sup> | *C. albicans* 14053<sup>a</sup> |
| Untreated | 0 | 8.71 ± 0.09 | 9.05 ± 0.02 | 8.24 ± 0.09 |
| Saline | 60 | 7.33 ± 0.08 | 7.95 ± 0.04 | 7.97 ± 0.07 |
| M-EDTA-25E | 15 | 5.62 ± 0.05 | 6.24 ± 0.15 | 7.44 ± 0.13 |
| | 30 | 5.48 ± 0.02 | 6.14 ± 0.14 | 6.66 ± 0.28 |
| | 45 | <1 | 6.18 ± 0.10 | 4.88 ± 0.03 |
| | 60 | <1 | 4.39 ± 0.02 | 4.43 ± 0.04 |
| PAD | 15 | 3.13 ± 0.10 | 6.40 ± 0.07 | 6.76 ± 0.05 |
| | 30 | <1 | 6.27 ± 0.22 | 3.69 ± 0.09 |
| | 45 | <1 | <1 | 1.91 ± 0.16 |
|  | 60 | <1 | <1 | <1 |

<sup>a</sup> $P \leq 0.001$ when compared with control; $n = 3$ ; detection limit 1 $\log_{10}$ CFU/ml.

^a^ *P* ≤ 0.001 when compared with control; *n* = 3; detection limit 1 log_10_ CFU/ml.

**Table 2.** Survivors (A) and regrowth (B) of clinical isolates following PAD treatment

| Lock Solution | Lock time (min) | Survivors ($\log_{10}$ CFU/ml ± SD ) | | |
| --- | --- | --- | --- | --- |
|  |  | *MRSA* SO385<sup>c</sup> | *S. epidermidis* M0881<sup>c</sup> | *C. albicans* P57072<sup>c</sup> |
| Untreated | 0 | 8.51 ± 0.17 | 7.37 ± 0.13 | 6.64 ± 0.22 |
| Saline | 60 | 6.66 ± 0.07 | 5.10 ± 0.16 | 5.62 ± 0.12 |
| M-EDTA-25E | 30 | 5.59 ± 0.19 | 3.61 ± 0.12 | 4.93 ± 0.13 |
| | 60 | 2.93 ± 0.20 | 2.84 ± 0.27 | 4.34 ± 0.34 |
| PAD | 30 | 2.67 ± 0.16 | 2.47 ± 0.25 | 3.01 ± 0.33 |
|  | 60 | <1 | <1 | <1 |

(B)
| Lock Solution | Lock time (min) | Regrowth ($\log_{10}$ CFU/ml ± SD ) | | |
| --- | --- | --- | --- | --- |
|  |  | *MRSA* SO385<sup>c</sup> | *S. epidermidis* M0881<sup>c</sup> | *C. albicans* P57072<sup>c</sup> |
| Untreated | 0 | 9.35 ± 0.07 | 9.40 ± 0.13 | 7.45 ± 0.16 |
| Saline | 60 | 9.10 ± 0.22 | 8.82 ± 0.27 | 7.42 ± 0.12 |
| M-EDTA-25E | 30 | 8.93 ± 0.10 | 9.03 ± 0.15 | 7.25 ± 0.17 |
| | 60 | 8.83 ± 0.14 | 8.53 ± 0.08 | 7.21 ± 0.22 |
| PAD | 30 | 8.94 ± 0.16 | 7.84 ± 0.26 | 6.55 ± 0.24 |
|  | 60 | <1 | <1 | <1 |

<sup>c</sup> $P \leq 0.05$ when compared with control; $n = 3$ ; detection limit 1 $\log_{10}$ CFU/ml.

^c^ *P* ≤ 0.05 when compared with control; *n* = 3; detection limit 1 log_10_ CFU/ml.

A detailed comparison between PAD and M-EDTA-25E was made for eradication and regrowth of *C. albicans* 14053 with extended lock times up to 360 minutes. Complete regrowth inhibition of *C. albicans* biofilm required 360 min with M-EDTA-25E, while a lock time of 60 minutes was needed for PAD (Table 3). Efficacy with 60-minute PAD lock solution was confirmed against 48-hour biofilms tested against microbes listed in Table 1 (data not shown).

**Table 3.** PAD and M-EDTA-25E efficacy at different lock dwell times against *C. albicans*.

| Lock time<br>(min) | *C. albicans* 14053 (log<sub>10</sub> CFU/ml ± SD) |  |  |  |
| --- | --- | --- | --- | --- |
|  | M-EDTA-25E |  | PAD |  |
|  | Survivors<sup>a</sup> | Regrowth<sup>a</sup> | Survivors<sup>a</sup> | Regrowth<sup>a</sup> |
| 0 | 7.25 ± 0.23 | 8.13 ± 0.27 | 7.25 ± 0.23 | 8.13 ± 0.27 |
| 30 | 5.89 ± 0.25 | 8.03 ± 0.22 | 5.25 ± 0.13 | 6.87 ± 0.19 |
| 60 | 4.87 ± 0.11 | 6.60 ± 0.17 | < 1 | < 1 |
| 120 | 4.34 ± 0.13 | 6.20 ± 0.32 | < 1 | < 1 |
| 180 | 2.60 ± 0.28 | 5.90 ± 0.30 | < 1 | < 1 |
| 240 | < 1 | 4.30 ± 0.27 | < 1 | < 1 |
| 300 | < 1 | 2.99 ± 0.25 | < 1 | < 1 |
| 360 | < 1 | <1 | < 1 | < 1 |

<sup>a</sup> $P \leq 0.001$ when compared with control; $n = 3$ ; detection limit 1 log<sub>10</sub> CFU/ml.

^a^ *P* ≤ 0.001 when compared with control; *n* = 3; detection limit 1 log_10_CFU/ml.

To confirm effective biofilm dispersal, Figure 2A shows that 60-minute exposure to PAD removed adhesive materials from inoculated glass slides with MRSA BAA-1707 and *C. albicans* 14053. By contrast, M-EDTA-25E exposure for 60 minutes led to considerable biomass remaining on glass slides (Figure 2A). Since adherent matter of *P. aeruginosa* biofilms was most difficult to remove (Figure 2B), their dispersal by PAD and M-EDTA-25E was quantified by means of optical absorption at 590 nm of the biofilm-infected glass slides (20). PAD exposure for 60 minutes reduced the optical absorption to approximately 1% compared to approximately 41% achieved with M-EDTA-25E (Figure 2B).

**Figure 2.**
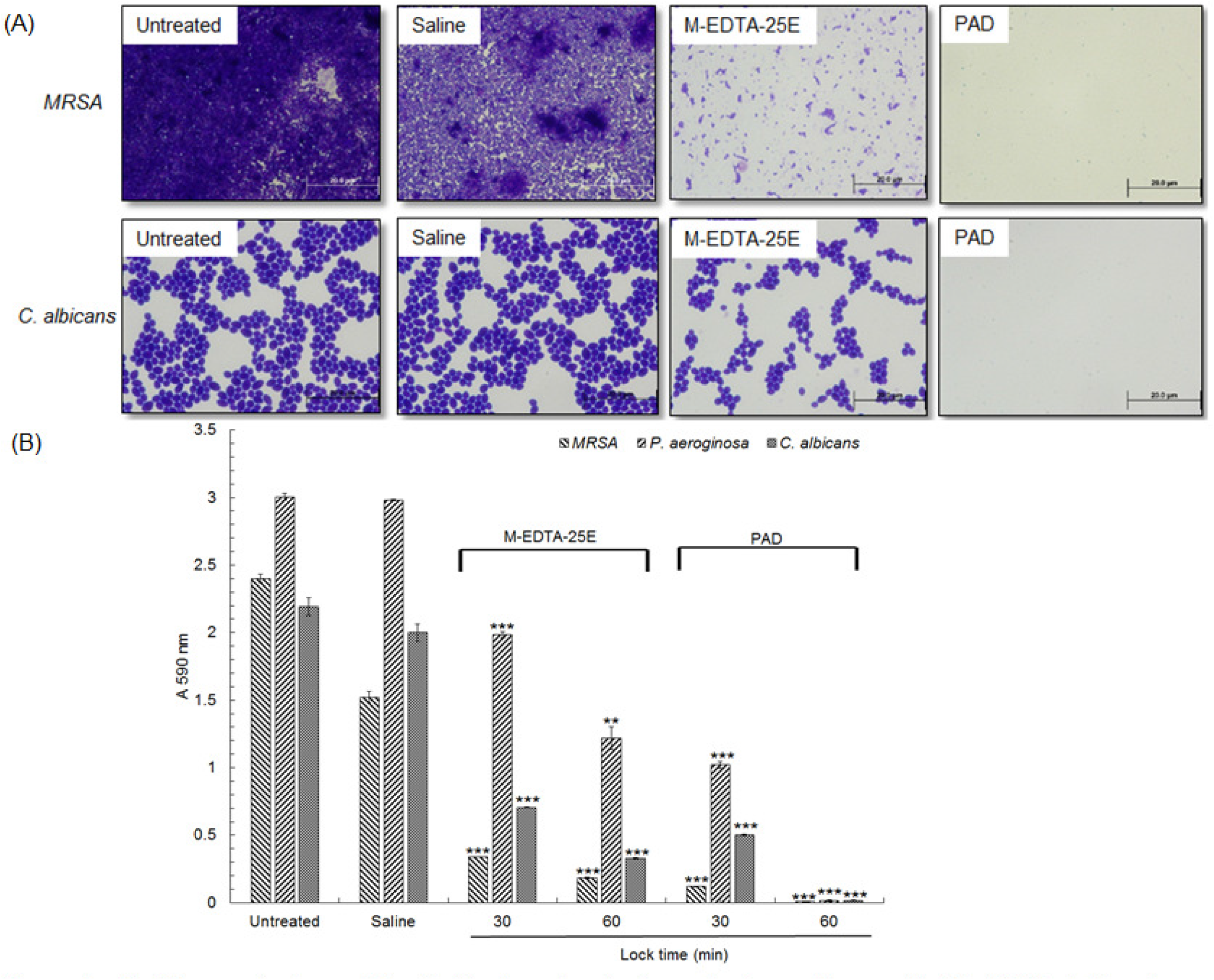
(A) Microscopic views of the biofilm formation in glass chambers after crystal violet (CV) staining (lens magnification, X 100) and (B) quantification of CV (B). \*\*\**P*≤ 0.005 and ** *P*≤ 0.01 when compared with control.

### Cytotoxicity to Primary Human Cells

Figure 3 shows that a simulated spill of PAD minimally affected the viability of HUVECs at all evaluation points of 0.5, 2, 15, and 24 hours. For the lock-to-medium ratio at both 0. 1% and 0.5%, there was no statistically significant difference in the viability between treated and untreated HUVECs at 24 hours, suggesting an effective repair of initial minor injuries in PAD-treated human cells. In contrast, the same amount of spilled M-EDTA-25E damaged HUVECs with cell viability clearly lower at the lock-to-medium ratio of 0.5% than 0.1%. Viability of HUVECs treated with M-EDTA-25E became progressively decreased with the incubation time, suggesting an accumulation of unrepaired injury. For the case of 24-hour incubation when necessary repairs of cellular damage should have taken place, cell viability was 59% for M-EDTA-25E at the lock-to-medium ratio of 0.5%, suggesting cell death.

**Figure 3.**
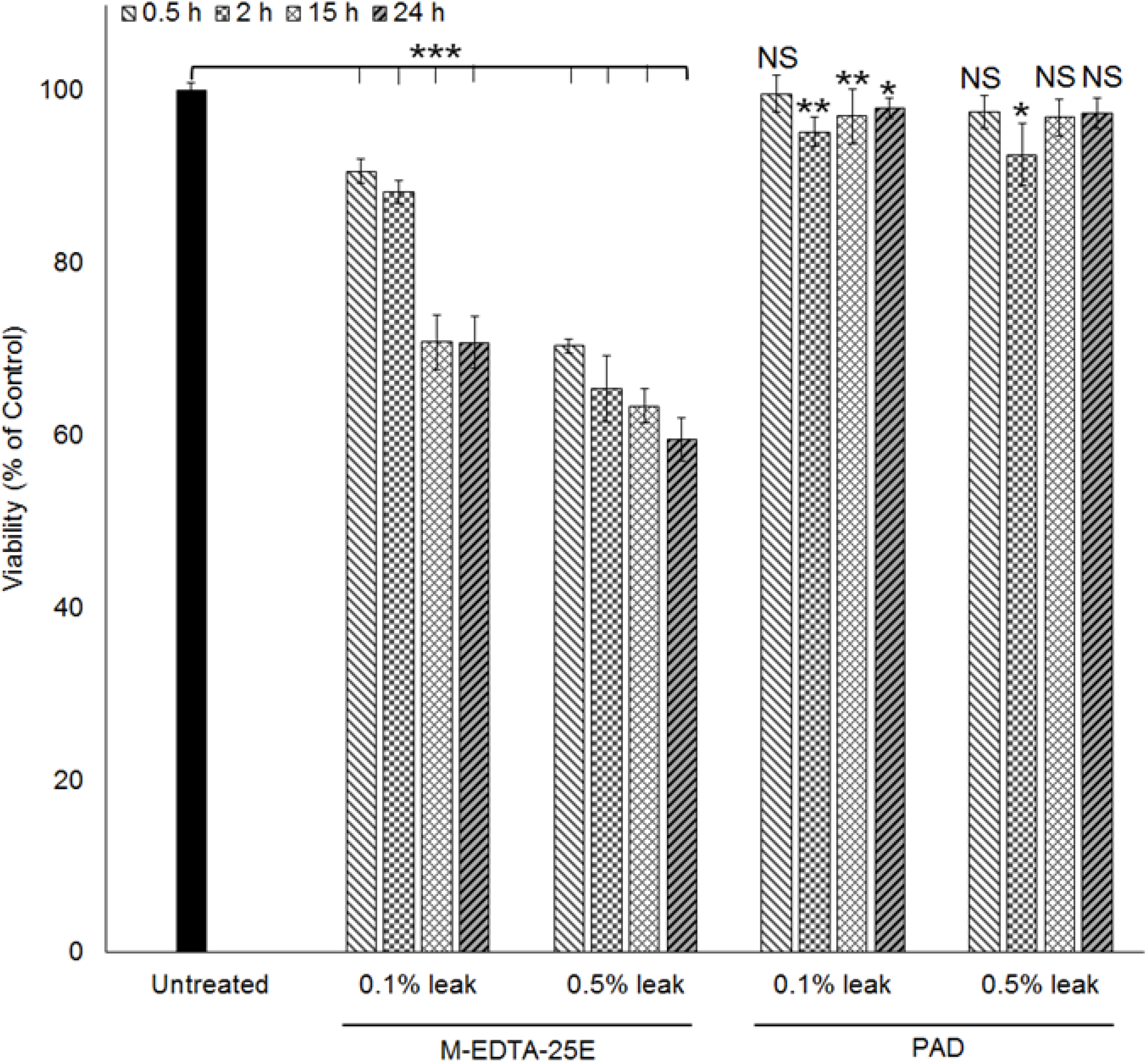
Evaluation of potential cytotoxic activity after exposure to HUVECs by leaked lock solution at a solution-to-medium ratio of 0.1% and 0.5%. *n* = 3. ** *P*≤ 0.01 and * *P* ≤ 0.05 when compared with the control.

### PAD effects on silicon catheter surface morphology

To test if the RONS in acidified PAD (Figure 1) impacts on catheter material after repeated PAD locking, we performed an accelerated aging test by continuously locking segments of catheter tubing for 7 days with regularly replenished PAD. In practice, PAD locking for 2 hours per day is sufficient to ensure complete suppression of microbial regrowth (Tables 1 and 2). There was no evidence of any episodes of fissures, cracks, or other morphological abnormalities in PAD-treated catheter tubes (Figure 4).

**Figure 4.**
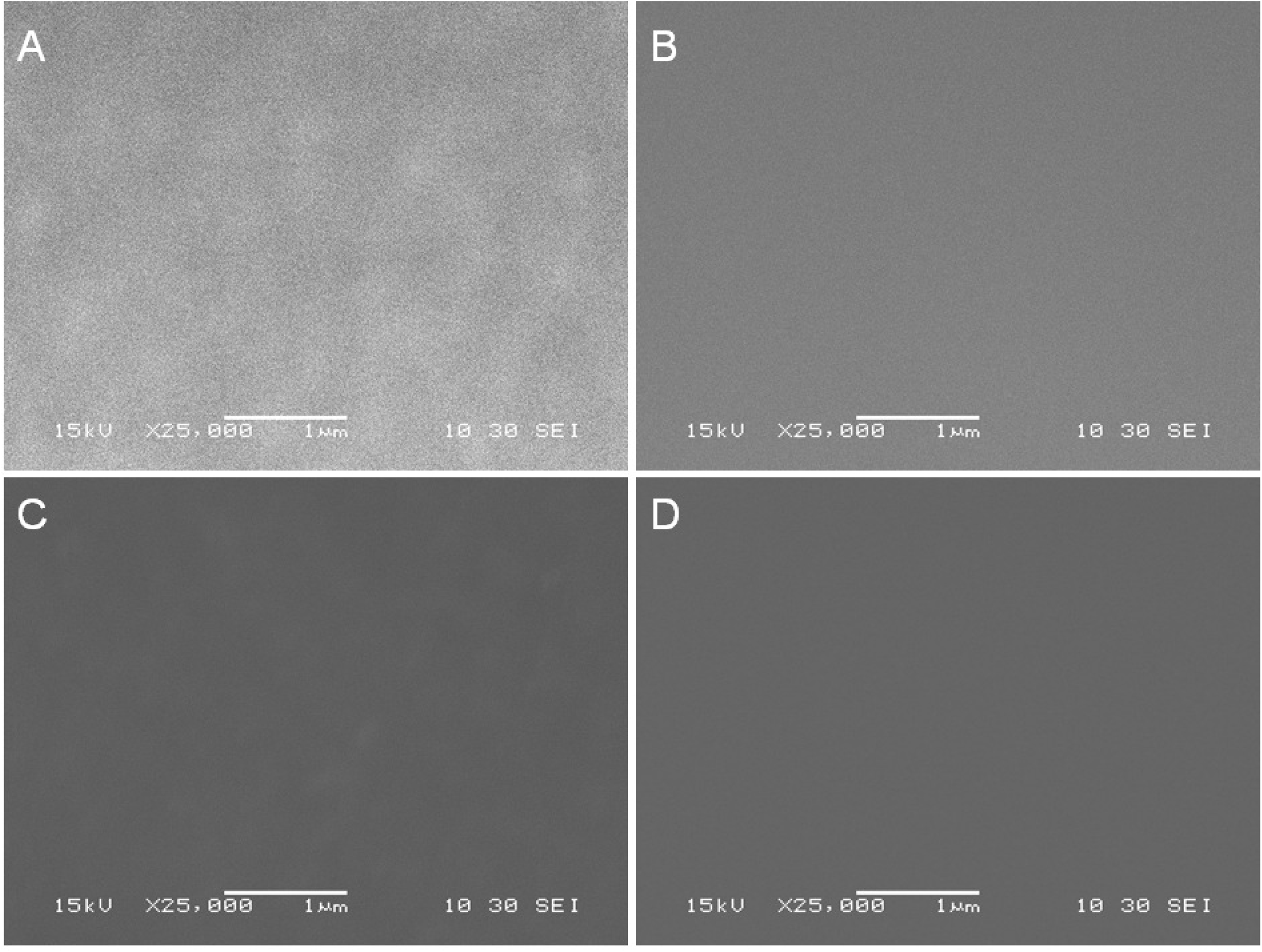
SEM micrographs (25000 X magnification) of the luminal surface of silicone catheter tubing segments with the tubing segments (A) as received from the manufacturer; (B) stored for 7 days at 37°C with no treatment; and treated with (C) saline or (D) PAD for 7 days at 37°C.

## DISCUSSION

Our study demonstrates that PAD lock solution is effective against microbial biofilms implicated in CRBSI with minimal cytotoxicity to primary human vein epithelial cells. Exposure to PAD lock for 60 minutes led to 6 - 8 order of magnitude reduction of viable bacterial and fungal biofilms and completely suppressed their regrowth. Similarly, PAD lock exposure for 60 minutes substantially reduced all adherent matter on glass slides inoculated with MRSA or *C. albicans*. For *C. albicans* 14053, M-EDTA-25E required 6 hours to achieve similar eradication and regrowth suppression. This is broadly in line with the capability of current ALT that generally requires prolonged lock times (29,30).

Variable efficacy has been reported for current ALT and antifungal lock therapy (AŕLT) (29,30). For PAD therapy, effective lock times were approximately 60 minutes against the Gram-positive and Gram-negative bacteria, and *C. albicans* we studied (Table 1, 2 and Figure 2). This suggests that PAD therapy can be applied prior to identification of pathogens that may be involved in a suspected CRBSI episode. The broad-spectrum antimicrobial properties of PAD are based on chemical diversity of reactive oxygen, nitrogen and chlorine species (31), and collectively capable of attacking different cellular targets (13,14) with minimal risk of systemic toxicity (Figure 3) or damage to silicone catheters (Figure 4).

An additional advantage of PAD is that we found minimal toxicity upon exposure to primary human umbilical vein endothelial cells. This is likely benefited from pulsed RONS of PAD (Figure 1) with which oxidative stresses to HUVECs are transient and as such are more easily repaired than chemicals that generate reactive oxygen species in living tissues for many months (32). Furthermore, continuous PAD locking for 7 days did not result in any morphological abnormalities of silicone such as fissures, cracks or damage when imaged using SEM (Figure 4). The maximum daily exposure of a CVC to PAD is 2 hours, and as such, the impact of continuous PAD locking for 7 days would be equivalent to that of a 2-hour daily PAD therapy on an indwelling CVC for 84 hospital days (=7×24/2).

Potential limitations to this study reflect the limited number of microbes included, studies were not carried out in plasma or serum, only silicone material was tested and additional testing was not performed to assess the impact of our PAD lock solution on the silicone tubes (e.g., testing the modulus of elasticity, force at break, and maximum stress at break).

PAD lock solution is free of antibiotics and the mechanism of action does not involve specific binding sites on bacteria or fungi, thereby minimizing the risk of selecting for antimicrobial resistance (11). In this context, it is of interest to note that a 2 hour exposure to a non-antibiotic containing, nitroglycerin-based lock solution reduces bacteria and fungal intraluminal biofilms by 4 orders of magnitude (19,34). Nitroglycerin produces reactive oxygen and nitrogen species (35) and this may explain why its activity against bacteria and fungi is similarly broad spectrum to PAD. Taken together, PAD lock therapy may be an innovative addition to current ALT and A*f*LT.

## ACKNOWLEDGEMENT

The authors declare no conflict of interest. HLC and MGK acknowledge support from Old Dominion University, USA.

## REFERENCES

1. Raad I, Hachem R, Hanna H, Bahna P, Charzinikolaou, Fang X, Jiang Y, Chemaly RF, Rolston K. 2007. Sources and outcome of bloodstream infections in cancer patients: the role of central venous catheters. Eur J Clin Microbiol Infect Dis 26: 549-556.

2. Worth LJ, Slavin MA. 2009. Bloodstream infections in haematology: Risks and new challenges for prevention. Blood Rev 23: 113-122.

3. Marschall J, Mermel LA, Fakih M, Hadaway L, Kallen A, O’Grady NP, Pettis AM, Rupp ME; Sandora T, Maragakis LL, Yokoe DS. 2014. Strategies to prevent central line-associated bloodstream infections in acute care hospitals. Infect Control Hosp Epidemiol. 35: 753-771.

4. Shah CB, Mitttelman MW, Costerton JW, Parenteau S, Pelak M, Arsenault R, Mermel LA. Antimicrobial activity of a novel catheter lock solution. 2002. Antimicrob Agent Chemother. 46: 1674-1679.

5. Ramos ER, Reitzel R, Jiang Y, Hachem RY, Chaftari AM, Chemaly RF, Hackett B, Pravinkumar SE, Nates J, Tarrand JJ, Raad II. 2011. Clinical effectiveness and risk of emerging resistance associated with prolonged use of antibiotic-impregnated catheters: More than 0.5 million catheter days and 7 years of clinical experience. Crit Care Med. 39: 245-251.

6. Niyyar VD, Lok CE. Pros and cons of catheter-lock solutions. 2013. Curr Opin Nephrol Hypertens. 22: 669-74.

7. Zakhour R, Chaftari AM, Raad I. 2016. Catheter-related infections in patients with haematological malignancies: Novel preventative and therapeutic strategies. Lancet Infect Dis; 16: e241-e250.

8. Mermel LA, Alang N. Adverse effects associated with ethanol catheter lock solutions: A systematic review. J Antimicrob Chemother 69: 2611–2619, 2014.

9. Luther MK, Bilida S, Mermel LA, LaPlante KL. Ethanol and isopropyl alcohol exposure increases biofilm formation in Staphylococcus aureus and Staphylococcus epidermidis. Infect Dis Ther 4:219-226, 2015.

10. Messing B, Peitra-Cohen S, Debure A, Beliah M, Bernier JJ. 1988. Antibiotic-lock technique: a new approach to optimal therapy for catheter-related sepsis in home-parenteral nutrition patients. J Parenter Enteral Nutr. 12: 185-189.

11. Spellberg B, Bartlett JG and Gilbert DN. 2013. The future of antibiotics and resistance. N Engl J Med. 368: 299-302.

12. Kong MG, Kroesen G, Morfill G, Nosenko T, Shimizu T, van Dijk J, Zimmermann JL. 2009. Plasma medicine: an introductory review. New J Phys. 11: 115012.

13. Vatansever F, de Melo WCMA, Avci P, Vecchio D, Sadasivam M, Gupta A, Chandran R, Karimi M, Parizotto NA, Yin R, Tegos GP, Hamblin MR. 2013. Antimicrobial strategies centered around reactive oxygen species - bactericidal antibiotics, photodynamic therapy, and beyond. FEMS Microbiol Rev. 37: 955-989.

14. Mai-Prochnow A, Murphy AB, McLean KM, Kong MG and Ostrikov K. 2014. Atmospheric pressure plasmas: Infection control and bacterial responses. Int J Antimicrob Agents. 43: 508-517.

15. Isbary G, Heinlin J, Shimizu T, Zimmermann JL, Morfill G, Schmidt HU, Monetti R, Steffes B, Bunk W, Li Y, Klaempfl T, Karrer S, Landthaler M, Stolz W. 2012. Successful and safe use of 2 min cold atmospheric argon plasma in chronic wounds: results of a randomized controlled trial. Brit J Dermatol. 167: 404-410.

16. Xu DH, Cui QJ, Xu YJ, Wang BC, Tian M, Li QS, Liu ZJ, Liu DX, Chen HL, Kong MG. Systemic study on the safety of immune-deficient nude mice treated by atmospheric plasma-activated water. Plasma Sci Technol 2018; 20: 044003.

17. Raad I, Henna H, Dvorak Tm Chaiban G, Hachem R. 2007. Optimal antimicrobial catheter lock solution, using different combinations of minocycline, EDTA, and 25-percent ethanol, rapidly eradicates organisms embedded in biofilms. Antimicrob Agent Chemother. 51: 78-83.

18. Chen C, Li F, Chen HL, Kong MG. 2017. Aqueous reactive species induced by a PCB surface microdischarge air plasma device: a quantitative study. J Phys D: App Phys. 50: 445208.

19. Reitzel RA, Rosenblatt J, Hirsh-Ginsberg C, Murray K, Chaftari AM, Hachem R, Raad I. 2016. *In vitro* assessment of the antimicrobial efficacy of optimized nitroglycerin-citrate-ethanol as a nonantibiotic, antimicrobial catheter lock solution for prevention of central line-associated bloodstream infection. Antimicrob Agent Chemother. 60: 5175-5181.

20. O’Toole GA. 2011. Microtiter dish biofilm formation assay. J Vis Exp. 30: 2437.

21. Polaschegg HD, Shah C. 2003. Overspill of catheter lock solution: safety and efficacy aspects. ASAIO J. 49: 713-715.

22. Polaschegg HD, Sodemann K. 2003. Risks related to catheter locking solutions containing concentrated citrate. Nephrol Dial Transplant. 18: 2688-2690.

23. Imlay JA, Linn S. Bimodal pattern of killing of DNA-repair-defective or anoxically grown *Escherichia coli* by hydrogen peroxide. 1986. J Bacteriol. 166: 519-527.

24. Lindemann C, Lupilova N, Muller A, Warscheid B, Meyer HE, Kuhlmann K, Eisenacher M, Leichert LI. 2013. Redox proteomics uncovers peroxynitrite-sensitive proteins that help *Escherichia coli* to overcome nitrosative stress. J Biol Chem. 288: 19688-19714.

25. Liu ZC, Guo L, Liu DX, Rong MZ, Chen HL, Kong MG. 2017. Chemical kinetics and reactive species in normal saline activated by a surface air discharge. Plasma Process Polym. 14: 1600113.

26. Wang L, Bassiri M, Najafi R, Najafi K, Yang J, Khosrovi. Hwong W, Barati E, Belisle B, Celeri C, Robson MC. 2007. Hypochlorous acid as a potential wound care agent. Part I. Stabilized hypochlorous acid: a component of the inorganic armamentarium of innate immunity. J. Burns Wounds. 6: 65-79.

27. Xia DS, liu T, Zhang CM, Yang SH, Wang SL. 2006. Antimicrobial effect of acidified nitrate and nitrite on six common oral pathogens *in vitro*. Chin Med J. 119: 1904-1909.

28. Clements WT, Lee S-R, Bloomer RJ. 2014. Nitrate ingestion: a review of the health and physical performances effects. Nutrients. 6: 5224-5264.

29. Vassallo M, Dunais B, Roger PM. 2015. Antimicrobial lock therapy in central-line associated bloodstream infections: a systemic review. Infection. 43: 389-398.

30. Walraven CJ, Lee SA. 2013. Antifungal lock therapy. Antimicrob Agent Chemother. 57: 1-8.

31. Carlsson S, Wiklund NP, Engstrand L, Weitzberg E, Lundberg JON. 2001. Effects of pH, nitric, and ascorbic acid on nonenzymatic nitric oxide generation and bacterial growth in urine. Nitric Oxide 5: 580-586.

32. Hu J, Liu Y-F, Wu C-F, Xu F, Shen Z-X, Zhu Y-M et al. 2009. Long-term efficacy and safety of all-trans retinoic acid/arsenic trioxide-based therapy in newly diagnosed acute promyelocytic leukemia. Proc Natl Acad Sci USA 106: 3342-3347.

33. Saivin S, Houin G. 1988. Clinical pharmacokinetics of doxycycline and minocycline. Clin Pharmacokinet. 15: 355-66.

34. Chaftari AM, Hachem R, Szvalb A, Taremi M, Granwehr B, Viola GM, Sapna A, Assaf A, Numan Y, Shah P, Gasitashvilli K, Nativdad E, Jiang Y, Slack R, Reizel R, Rosenblatt J, Mouhayer E, Raad I. 2017. A novel nonantibiotic nitroglycerin-based catheter lock solution for prevention of intraluminal central venous catheter infections in cancer patients. Antimicrob Agent Chemother. 61: e00091-17.

35. Wenzel P, Mollnau H, Oelze M, Schulz E, Wickramanayake JMD, Muller J, Schuhmacher S, Hortmann M, Baldus S, Gori T, Brandes RP, Munzel T, Daiber A. 2008. First evidence for a crosstalk between mitochondrial and NADPH oxidase-derived reactive oxygen species in nitroglycerin-triggered vascular dysfunction. Antioxid Redox Signal. 10: 1435-1447.

